# *Cryptococcus neoformans* secretes small molecules that inhibit IL-1β inflammasome-dependent secretion

**DOI:** 10.1101/554048

**Authors:** Pedro Henrique Bürgel, Clara Luna Marina, Pedro H. V. Saavedra, Patrícia Albuquerque, Paulo Henrique Holanda, Raffael de Araújo Castro, Heino Heyman, Carolina Coelho, Radames J. B. Cordero, Arturo Casadevall, Joshua Nosanchuk, Ernesto Nakayasu, Robin C. May, Aldo Henrique Tavares, Anamelia Lorenzetti Bocca

## Abstract

*Cryptococcus neoformans* is an encapsulated yeast that causes disease mainly in immunosuppressed hosts. It is considered a facultative intracellular pathogen because of its capacity to survive and replicate inside phagocytes, especially macrophages. This capacity is heavily dependent on various virulence factors, particularly the glucuronoxylomannan (GXM) component of the polysaccharide capsule, that render the non- or poorly-activated macrophage ineffective against phagocytosed yeast. Strategies utilized by macrophages to prevent this scenario include pyroptosis (a rapid highly inflammatory cell death) and vomocytosis (the expulsion of the pathogen from the intracellular environment without lysis). Inflammasome activation in phagocytes is usually protective against fungal infections, including cryptococcosis. Nevertheless, recognition of *C. neoformans* by inflammasome receptors requires specific changes in morphology or the opsonization of the yeast, impairing a proper inflammasome function. In this context, we analyzed the impact of molecules secreted by *C. neoformans* B3501 strain and its acapsular mutant *Δcap67* in an inflammasome activation *in vitro* model. Our results showed that conditioned media derived from B3501 was capable of inhibiting inflammasome dependent events (i. e. IL-1β secretion and LDH release via pyroptosis) more strongly than conditioned media from *Δcap67*, regardless of GXM presence. We also demonstrated that macrophages treated with conditioned media were less responsive against infection with the virulent strain H99, exhibiting lower rates of phagocytosis, increased fungal burdens and enhanced vomocytosis. Moreover, we showed that the aromatic metabolite DL-Indole-3-lactic acid (ILA) was present in B3501’s conditioned media and that this fungal metabolite is involved in the regulation of inflammasome activation by *C. neoformans*. Overall, the results presented show that conditioned media from a wild-type strain can inhibit an important recognition pathway and subsequent fungicidal functions of macrophages, contributing to fungal survival *in vitro* and suggesting that this serves as an important role for secreted molecules during cryptococcal infections.

**Author’s Summary:** *Cryptococcus neoformans* is the agent of cryptococcal meningitis, a disease that can be life-threatening in immunocompromised hosts such as those infected with HIV. The infection thrives in hosts that poorly activate their immune system, mainly because of the yeast’s ability to survive inside macrophages and migrate towards the central nervous system. Emerging data indicate that cryptococci modulate the host immune response, but the underlying mechanisms remain largely uncharacterized. Here we show that secreted molecules from a wild-type strain of *C. neoformans* impair inflammatory responses driven by inflammasome activation, which in turn impact the macrophage antifungal activity. We further show that this inhibition does not involve GXM, the main constituent of the fungal capsule, but rather is partially dependent on DL-Indole-3-lactic acid (ILA), a metabolite not previously implicated in fungal virulence.

## Introduction

*Cryptococcus neoformans* is a fungal pathogen that primarily affects immunocompromised patients with acquired immunodeficiency syndrome (AIDS) [1]. *C. neoformans* is responsible for over 180 thousand deaths yearly worldwide [2]. Human infection is usually initiated by the inhalation of environmental spores or yeasts that are frequently present in avian droppings [3–7]. Once in the lung, the fungus is usually cleared from the host or survives within granulomas. In the context of immunosuppression, primary acquisition or relapse of previously contained yeast can result in disseminated disease, especially involving the central nervous system [5,8,9].

The ability of *C. neoformans* to remain viable and survive inside the host is dependent on its many virulence factors, which allow the fungus to modulate and evade the immune response [10,11]. These virulence factors include enzymes (laccase, urease, phospholipases, proteases and others) that can be secreted freely or encapsulated in extracellular vesicles [10,12–14], melanin deposition in the cell wall [14,15] and the formation of a polysaccharide, which is considered the most important of these factors [16–18]. Glucuronoxylomannan (GXM) is the most prevalent of these capsular polysaccharides, facilitating *C. neoformans* resistance against phagocytosis and suppressing various immune responses [19–26]. Altogether these virulence factors enable this fungus to effectively survive and thrive as a facultative intracellular pathogen, particularly within macrophages [27–33]. In fact, depletion of alveolar macrophages in mice is associated with a worse prognosis, indicating that they are coopted by *C. neoformans* during pathogenesis to facilitate fungal growth and dissemination [34]. Phagocytic cells that are unable to kill intracellular yeast cells can return fungal cells to the extracellular environment, either through a non-lytic exocytosis called vomocytosis [35–38] or a rapid, highly inflammatory and inflammasome dependent cell death referred to as pyroptosis [39–41].

The inflammasome is an intracellular multi-protein complex that usually requires an intracellular damage-associated molecular pattern (DAMP) for its oligomerization and proper function [42]. The canonical activation step requires the engagement of an intracellular receptor from the NOD-like receptors (NLRs) or AIM2-like receptors (ALRs) family, an adaptor protein ASC and the cleavage of procaspase-1. Although some cell types are able to activate inflammasome pathways from basal expression levels, most of them require extracellular signaling, promoted by membrane bound pattern-recognition receptors, to initiate inflammasome activation [43]. The activated caspases in this context are responsible for the previously described pyroptosis cell death and, most importantly, for the processing of interleukin (IL)-1β and IL-18, important mediators of inflammatory Th1/Th17 driven responses [44].

Among the receptors associated with inflammasome oligomerization, NLRP3 is one of the best described and well-characterized in fungal recognition [45]. This receptor is involved in the recognition of various fungal species, between yeast and hyphal forms, and opportunistic and primary pathogens [46–51]. The activation of NLRP3 is usually dependent on one or more intracellular stress signals (i. e. potassium efflux; mitochondrial ROS production and cathepsin release), which are associated with the interaction between the host cell and the fungus [42,45]. NLRP3 activation in response to *C. neoformans* only occurs when the yeast is in specific conditions such as biofilms [52], lacking capsule [53], or opsonized prior to phagocytosis [54]. Moreover, all three classical stress signals are required to activate NLRP3 during these interactions [52]. Notably, mice lacking NLRP3 or ASC are more susceptible to cryptococcal infection with encapsulated yeast cells, whereas infection with acapsular yeast cells results in higher fungal burdens in the lungs in NLRP3 knockout mice [52,53]. Likewise, susceptibility to cryptococcal infection has been also observed in murine knockout models for IL-1β and IL-18 receptors [55,56].

Different strains of *C. neoformans* elicit variable IL-1β induction, especially in *in vitro* models. Although GXM participates in inflammasome inhibition when macrophages are challenged with acapsular strains [53], capsule-independent inhibition of the inflammasome remains poorly understood. Here we show that other secreted molecules besides GXM can specifically interfere with intracellular signals during inflammasome activation, suppressing various processes associated with this activation and reducing the overall antifungal capacity of macrophages. Furthermore, we have defined one molecule present in *C. neoformans* conditioned media that participates in inhibiting inflammasome activation in the presence of this remarkable fungus.

## Results

### Exopolysaccharide incorporation on acapsular mutant does not impair macrophage inflammasome activation

As described [53,54], *C. neoformans* mutants lacking GXM expression and proper capsule formation triggered inflammasome activation more effectively than compared to wild type encapsulated yeast cells (Fig 1A). However, we found that *Δcap67* encapsulated with exo-PS induced significantly more IL-1β secretion compared to the wild type B3501 (Fig 1B). This result suggested that the presence of GXM on the yeast surface by itself was not sufficient to explain the differences in inflammasome activation observed between acapsular mutants and their wild type counterparts.

**Fig 1.**
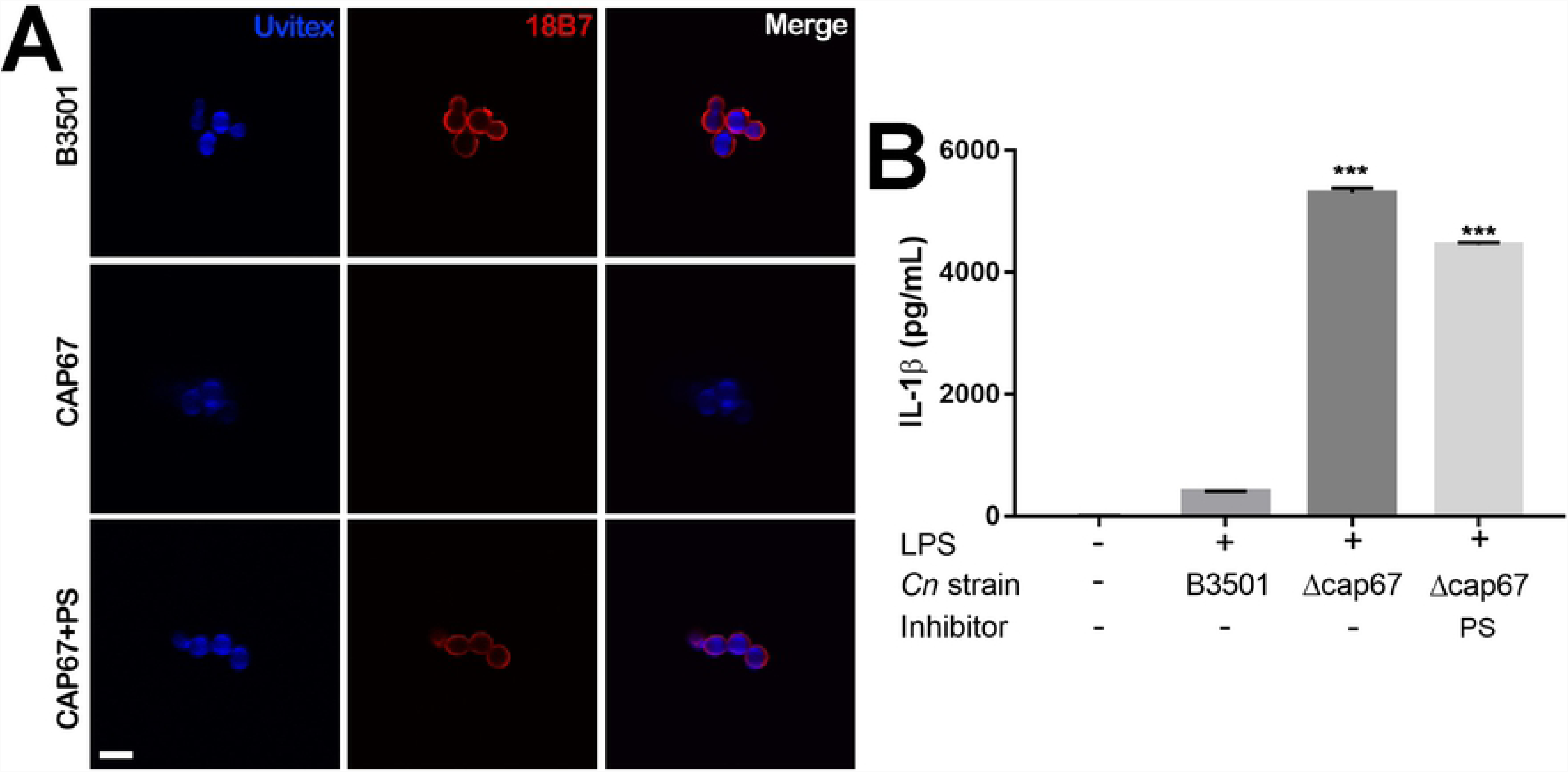
Exopolysaccharide incorporation on an acapsular mutant does not impair macrophage inflammasome activation. A. GXM detection by immunofluorescence of B3501, CAP67 (*Δcap67*) and CAP67+PS (*Δcap67* coated with polysaccharides from B3501) labeled with Uvitex2B (blue) and mAb against GXM (18B7) (red). B. BMMs were stimulated with LPS (500ng/mL) and infected with opsonized B3501, *Δcap67* or *Δcap67*+PS strains (MOI 2:1) for 24 h. Statistical analysis was performed using One-way ANOVA, where ***: P ≤ 0.001. Comparisons were made with the B3501 infected group.

### CM35, but not CMCAP or minimal media, reduces IL-1β secretion

Since many of the virulence factors presented by *C. neoformans* are secreted [10,15], we evaluated whether components released by the fungus were able to inhibit inflammasome canonical activation, induced by the combination of LPS and nigericin treatments (“activated” macrophages and dendritic cells). The addition of CM35 to activated macrophages and dendritic cells significantly reduced the secretion of IL-1β by these cells, while CMCAP, MM and glycine inhibited IL-1 β secretion to a lesser degree (Fig 2A and 2B, S1A). Interestingly, CM35, CMCAP and MM did not reduce the secretion of TNF-α (Fig 2C and 2D, S1B), even if added before LPS priming (S1C Fig). The specific interference of conditioned media in the secretion of IL-1β, an inflammasome dependent cytokine, but not in TNF-α, an inflammasome-independent cytokine, indicates that the canonical inflammasome pathway is being inhibited. The reduction in IL-1β observed with CMCAP and minimal media is likely explained by the presence of glycine, since this component is present in minimal medium and exhibits the same interference in the cytokine secretion (Fig 2A). When primed with LPS and challenged with the acapsular strain *Δcap67*, macrophages still show a marked reduction in IL-1β levels in the presence of CM35, while in these conditions MM and glycine did not affect the secretion of this cytokine (Fig 2E), indicating that CM35 is able to affect the inflammasome pathway in the context of different activators. To investigate the molecular identity of the component causing effects in the inflammasome, the conditioned and minimal medium were filtered to obtain fractions of different sizes. Of the fractions from CM35, all fractions, except the fraction between 100 and 10 kDa inhibited IL-1β secretion (Fig 2F), indicating that molecules below the 10 kDa range were necessary for the effect seen with CM35 treatment. Since the greatest effect was achieved with the fractions below 1kDa, the next assays were conducted utilizing this small size fraction (1KDa-CM35), unless stated otherwise.

**Fig 2.**
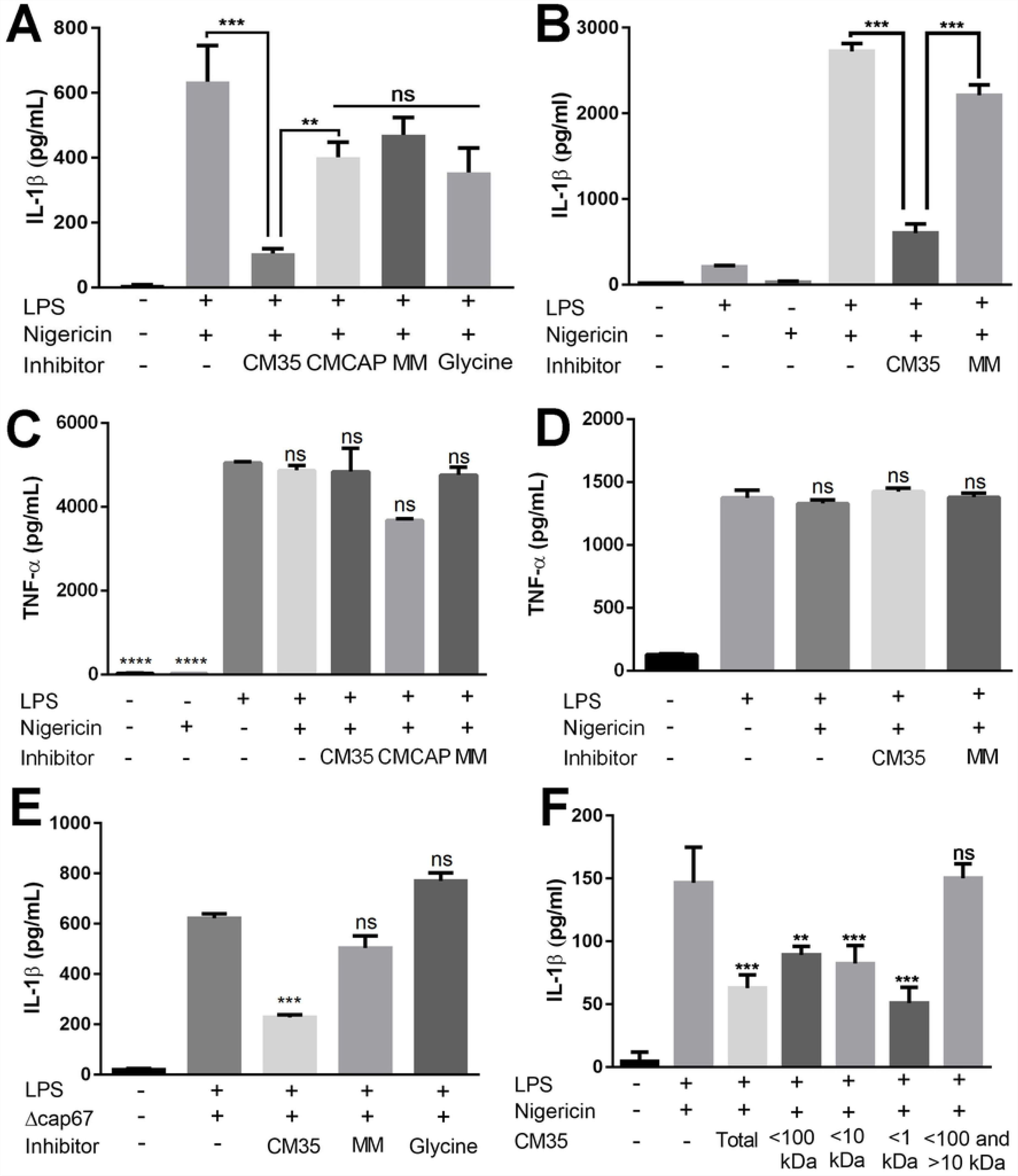
CM35, but not CMCAP or Minimal Media reduces IL-1β secretion. A,C. BMMs were stimulated with LPS (500ng/mL) and/or nigericin (20µM), added or not by mediums (CM35; CMCAP or MM 10% v/v) or glycine (13mM) overnight (18h). B,D. DCs were stimulated with LPS (500ng/mL) and/or nigericin (20µM), added by media (CM35; CMCAP or MM 10% v/v) or glycine (13mM) overnight (18h). E. BMMs were stimulated with LPS (500ng/mL), infected with *Δcap67* strain (MOI 5:1), added or not by mediums (CM35 or MM 10% v/v) or glycine (13mM) overnight (18h). F. BMMs stimulated with LPS (500ng/mL) and/or nigericin (20µM), added by CM35>1kDa (10% v/v) overnight (18h). Supernatants were collected after stimulus and cytokines were quantified by ELISA technique. CM35 = Conditioned Media from B3501; CMCAP = Conditioned Media from *Δcap67*; MM = Minimal Media. Statistical analysis was performed utilizing One-way ANOVA, where ns: not significant; **: P ≤ 0.002; ***: P ≤ 0.001. Comparisons were made with the positive control group (LPS + Nigericin) when not indicated (C-F).

### CM35 inhibits caspase-1 activation, promoting pro-IL-1β accumulation and cell death inhibition

Canonical inflammasome activation triggers various processes in the cell, including but not limited to IL-1β maturation and secretion. In that context, we analyzed other cell processes related to inflammasome activation that might be altered in macrophages interacting with CM35, while primed with LPS and nigericin, such as cell death via pyroptosis and intracellular pro-IL-1β cleavage. The results of the measurements of cell death by LDH release followed the same pattern seen in IL-1β secretion (Fig 3A). The fact that both stimuli, LPS and nigericin, were necessary for LDH release confirms cell death via inflammasome activation in our experimental conditions. When macrophages were treated with CM35, LDH release was largely abrogated, while treatment with CMCAP or MM resulted in a smaller inhibition of macrophage cell death (Fig 3A), which again is likely partly explained by the presence of glycine in the media [57]. Regarding pro-IL-1β, the results showed that macrophages must be primed with LPS for this cytokine production and that the presence of nigericin does not alter its intracellular levels, indicating a constant production of this inactive cytokine in our model. (Fig 3B). When treated with CM35, primed macrophages exhibited an increase in intracellular pro-IL-1β protein, while cells treated with MM did not (Fig 3B). Surprisingly, treatment with CMCAP also induced a high accumulation of intracellular pro-IL-1β (Fig 3B). This might be explained by the fact that CMCAP was the only treatment able to induce pro-IL-1β production without LPS. In fact, both conditioned media increased *IL1β* (i. e. pro-IL-1β) transcription levels in primed macrophages, while not similarly inducing *nfκb* transcription, even after 24 h after LPS stimuli (S2 Fig). CMCAP alone was also able to induce TNF-α secretion by unprimed macrophages (Fig 3C), corroborating the assumption that this conditioned media can induce first signaling via NF-κB activation.

**Fig 3.**
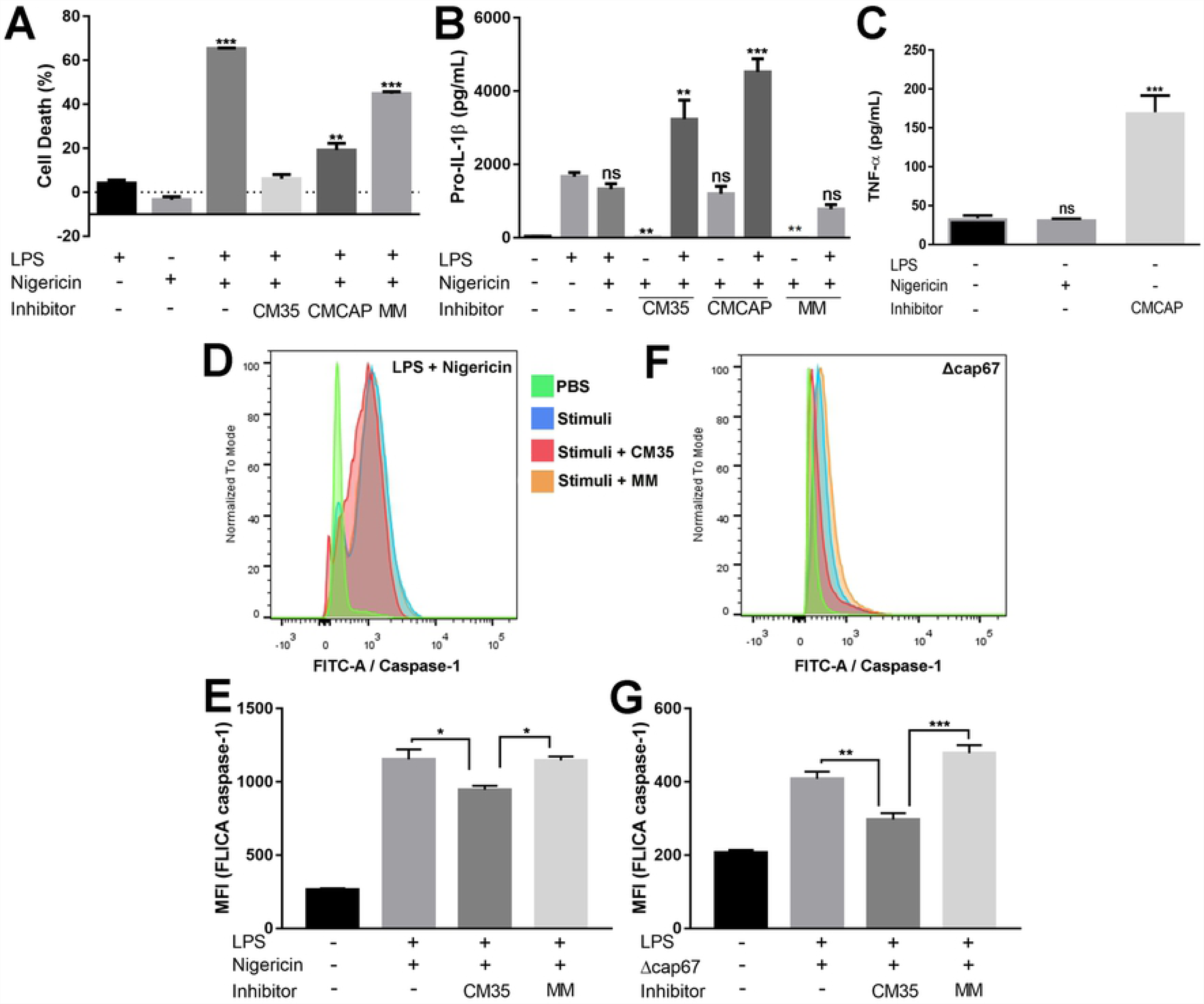
CM35 inhibits caspase-1 activation, promotes pro-IL-1β accumulation and inhibits cell death. A. LDH release from supernatants of BMMs stimulated with LPS (500ng/mL) and/or nigericin (20µM), with or without inhibitors (10% v/v). Medium group was utilized as negative control (0% of cell death) and DMSO (15%) group was used as positive control (100% of cell death) B. Pro-IL-1β production measured from cell lysates of BMMs stimulated with LPS (500ng/mL) and/or nigericin (20µM), with or without inhibitors (10% v/v). C. IL-1β release measured from supernatants of BMMs stimulated with nigericin (20µM), with or without CMCAP. D-G. BMMs stimulated with LPS (500ng/mL) and nigericin (20µM) or *Δcap67* (MOI 5:1), with or without inhibitors were analyzed for caspase-1 activation (FLICA). LDH release was measured by colorimetric assay (A), cytokines were measured by ELISA (B,C), and caspase activation was measured by flow cytometry (D-G). Statistical analysis was performed by One-way ANOVA, where ns: not significant; *: P ≤ 0.033; **: P ≤ 0.002; ***: P ≤ 0.001. Comparisons were made with the LPS group (A,B), medium group (C) and positive group (LPS + Nigericin) (D).

Having confirmed that CM35 inhibited various signals related to inflammasome activation, the next step was to observe inflammasome activation itself, analyzing the last step in the multi-protein complex assembly: the auto-cleavage of pro-caspase-1 into caspase-1 active form. The measurement was done with single cell analysis, using flow cytometry (Fig 3D-3G). Results showed that primed macrophages treated with CM35 decrease the caspase-1 active form, compared to untreated cells or cells treated with MM. Moreover, this was independent of the second signal activator used: nigericin (Fig 3D and 3E) or infection with *C. neoformans* strains (Fig 3F and 3G). These results are consistent with the finding that CM35 inhibits IL-1β secretion, showing that CM35 can inhibit inflammasome activation induced via different stimuli (i.e. nigericin and infection with *Δcap67*). Altogether, these results indicate that CM35 interfered with inflammasome activation, affecting the later stages including caspase-1 activation, pro-IL-1β cleavage, IL-1β secretion and pyroptotic cell death.

### CM35 impacts phagocytic capacity and extrusion events in interactions between macrophages and *C. neoformans*

A potent pro-inflammatory polarization is important for a protective response against cryptococcosis [58]. Given that IL-1β is one of the key cytokines associated with heightened inflammation, we analyzed if CM35 treatment caused a functional impairment of macrophages when infected with *C. neoformans*. We initially evaluated intracellular fungal burdens in macrophages when yeast cells were added at the same time as conditioned media or minimal media. Interestingly, none of the treatments impacted fungal intracellular growth in primed macrophages (Fig 4A). However, when macrophages were primed for 18 h with conditioned medium, CM35 promoted a significant reduction in the phagocytic capacity presented by macrophages, with or without LPS stimuli (Fig 4B).

**Fig 4.**
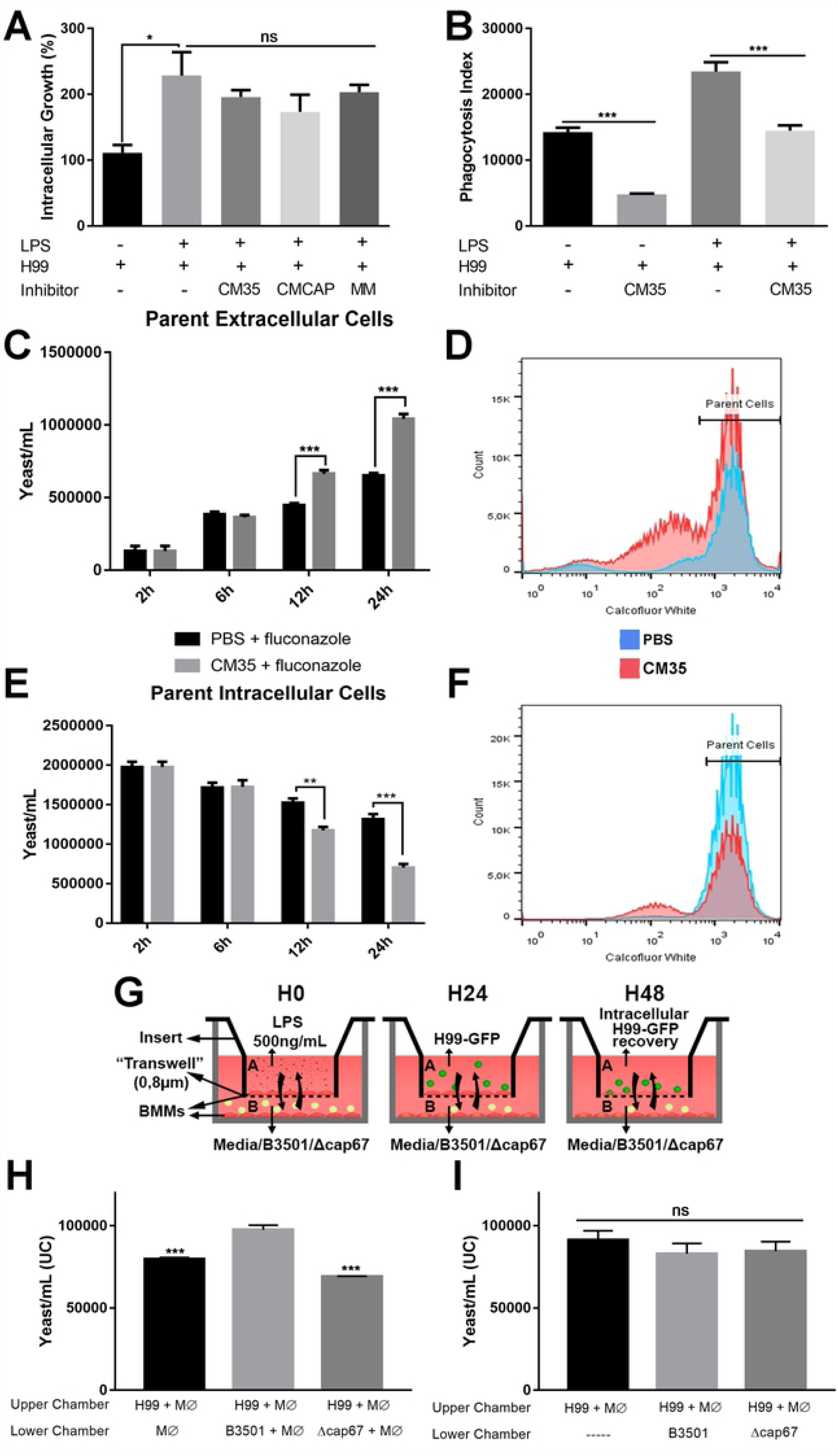
CM35 impacts phagocytic capacity and extrusion events in interactions between macrophages and *C. neoformans*. A. Intracellular growth (CFU of 3h vs 24h post infection) of yeasts cells in BMMs stimulated with LPS (500ng/mL) and H99 (2:1), with or without inhibitors (10% v/v). B. Phagocytosis index (2h post infection) from BMMs stimulated with LPS (500ng/mL) and H99 (5:1), with or without CM35 (10% v/v). C-F. Flow cytometry analysis of extracellular (C and D) and intracellular (E and F) yeast cells (Calcofluor white high – 2h, 6h, 12h and 24h post infection) from BMMs infected with H99 (10:1), with or without CM35 (10% v/v). I. Scheme for transwell infection assay, illustrating the steps taken during the assay. H-I. Flow cytometry measurement (24h post infection) of intracellular yeast cells in BMMs infected with H99 (2:1), in the upper chamber of a transwell apparatus. In the lower chamber, BMMs were stimulated with LPS (500ng/mL) and B3501 or *Δcap67* (5:1) (G). Alternatively, the lower chamber contained only yeast cells from B3501 or *Δcap67* in a media with LPS (500ng/mL) (H). Statistical analysis was performed utilizing One-way ANOVA, where ns: not significant; *: P ≤ 0.033; **: P ≤ 0.002; ***: P ≤ 0.001. Comparisons were made with the B3501 lower chamber infected group (G)

Another important aspect of macrophage-*Cryptococcus* interaction that we analyzed was vomocytosis in the presence of CM35. Parental H99 cells were labelled with calcofluor white to differentiate parent from daughter yeast cells (daughter cells form de novo cell walls and are thus not labeled). Macrophages were concomitantly exposed to CM35 and infected with H99 strain. Yeast cells with high fluorescence were measured to exclude cells that appeared via budding. We observed that groups containing macrophages treated with CM35 had a higher content of extracellular parent yeast cells compared to the groups containing untreated macrophages (Fig 4D and 4F). The difference between groups starts to increase at 12 h of infection (p>0.001), increasing once again after 24 h. Corroborating this data, intracellular parent yeast cells number showed an inverse pattern, reducing after 12 h of infection in the groups treated with CM35 (Fig 4E and 4G). Knowing that macrophages treated with CM35 present an equal rate of cell death when compared to untreated cells (Fig 3A), the increase of extracellular cells observed in this assay likely indicates an increase in vomocytosis rate in the presence of CM35.

To further study the impact of secreted molecules in macrophage function, a transwell assay involving two sequential infections in different chambers physically separated by a 0.8 µm membrane was carried out (Fig 4G). Flow cytometry assays were performed to confirm that the yeast cells present in one chamber would not cross to the other chamber. H99 cells expressing GFP were used for the infection of the macrophages in the upper chamber, while non-fluorescent B3501 or *Δcap67* were used for infection in the lower chamber. After 24 h, the intracellular yeasts from these macrophages were recovered after lysis. All yeast cells recovered from the upper chamber exhibited high GFP fluorescence, indicating no crossing from yeast cells present in the lower chamber (S3 Fig). A significantly higher intracellular fungal burden was observed in macrophages infected with H99 strain in a chamber vertically adjacent to the bottom chamber containing macrophages infected with B3501 strain compared to H99 growth when the adjacent chamber contained macrophages alone or macrophages infected with *Δcap67* strain (Fig 4H). Interestingly, this increase in fungal burden derived by B3501 was not observed when only yeast cells were seeded in the bottom chambers, with macrophages present just in the upper chambers (Fig 4I).

Overall, these experiments indicate that CM35 alters macrophage function at late timepoints, potentially by inhibiting a pro-inflammatory environment.

### CM35 inhibition characteristics indicate that a small and polar, non-polysaccharide, molecule is responsible for the effects

To further characterize the conditioned medium, CM35 was submitted to autoclaving and separated into polar and nonpolar soluble fractions. Autoclaved CM35 retained its inhibitory properties and the polar fraction, but not the nonpolar, also retained this property (Fig 5A). Raw CM35 was also treated with various proteases before its addition to primed macrophages. None of the treatments eliminated the inhibition promoted by the conditioned media, and the proteases themselves also did not significantly alter IL-1β secretion (Fig 5B). Overall, the analysis showed that a small, polar, and heat and protease resistant molecule (or molecules) resulted in the CM35 inhibitory properties.

**Fig 5.**
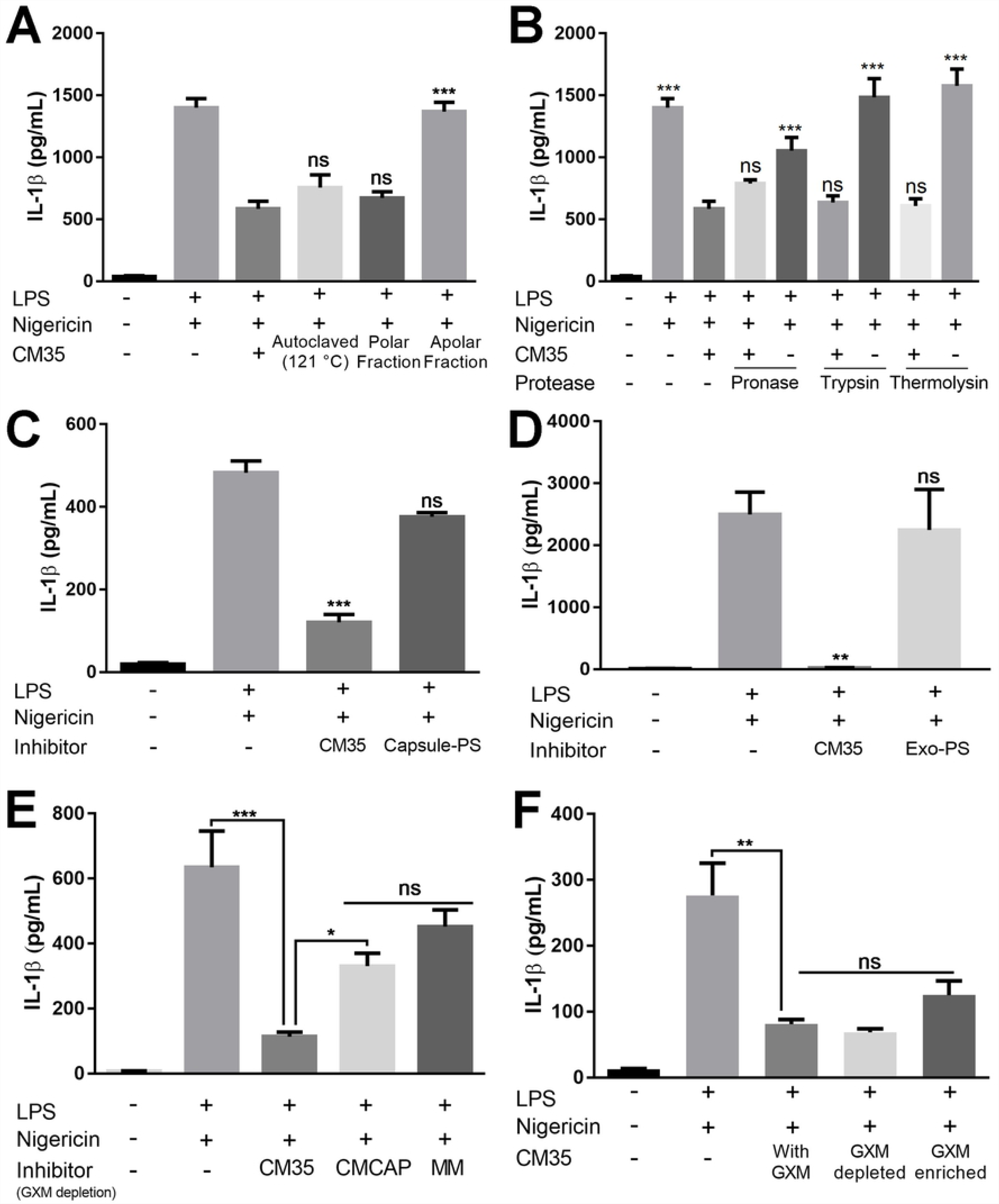
CM35 inhibition characteristics indicate a small, polar molecule that does not derive from a polysaccharide origin. A-B. BMMs were stimulated with LPS (500ng/mL) and/or nigericin (20µM), with or without CM35 fractioned by water affinity, autoclaved (B) or treated with proteases (C) (10% v/v) overnight (18h). C-D. BMMs were stimulated with LPS (500ng/mL) and/or nigericin (20µM), with or without potential inhibitor (10% v/v) overnight (18h). Polysaccharides (200ug/mL) extracted from capsule (C) or secreted in minimal medium (D) were used as inhibitors. E-F. BMMs were stimulated with LPS (500ng/mL) and/or nigericin (20µM), with or without possible inhibitor (10% v/v) overnight (18h). Conditioned medium and minimal media (CM35, CMCAP and MM) were treated for GXM depletion utilizing capture ELISA and used as inhibitors (10% v/v) (E). GXM recovered from ELISA was mixed with CM35 and utilized as inhibitor (10% v/v) (F). Supernatants were recovered after stimuli and IL-1β secretion was quantified using ELISA. Statistical analysis was performed by One-way ANOVA, where ns: not significant; *: P ≤ 0.033; **: P ≤ 0.002; ***: P ≤ 0.001. Comparisons were made with the CM35 treated group (A-B) or the positive control group (LPS + nigericin) (C-D).

GXM is a known virulence factor that matches with the characteristics possessed by our candidate molecule. While generally depicted as a high molecular weight polysaccharide, derived GXM molecules can be detect within smaller sizes [59]. We decided to investigate the role of GXM and GXM-derived molecules when interacting with activated macrophages and in the context of CM35 treatment, i.e., if we could detect these molecules in macrophages treated with CM35 below 1kDa size fraction, but not if treated with CMCAP (S4 A-C Fig). Firstly, polysaccharides derived from the yeast capsule (Fig 5C) or exo-polysaccharides obtained from the culture media (Fig 5D) were used to treat macrophages primed with LPS and nigericin. None of these treatments were able to significantly reduce IL-1β secretion compared to CM35, suggesting that GXM is not enough to promote inflammasome inhibition. Similarly, depleting GXM from CM35 using mAb 18B7 capture protocol (S2D-E Fig) [60] did not alter IL-1β secretion by activated macrophages (Fig 5E). Supporting the hypothesis that GXM is not the candidate molecule, the recovery of GXM from the coated ELISA plate and enrichment of its content in the CM35 did not significantly affect IL-1β secretion (Fig S2F and 5F). Although GXM has the properties exhibited by our candidate molecule, these assays indicate that GXM may not have an important role on inflammasome inhibition seen in our model.

### DL-Indole-3-lactic acid (ILA) participates in inflammasome inhibition property possessed by CM35

Given that GXM did not interfere in our inflammasome activation model, we pursued the identification of our active candidate molecule by analyzing CM35 and CMCAP using mass spectrometry. Three candidates met our criteria of being small, polar, and heat and protease resistant molecules (Fig 5): DL-3-Phenyllactic acid (PLA); DL-p-Hydroxyphenyllactic acid (HPLA) and DL-Indole-3-lactic acid (ILA) (S4 Fig). All three are aromatic metabolites derived from amino acid metabolism and are produced by several species of prokaryotes and eukaryotes [61,62]. Testing these three metabolites individually revealed that ILA was the most inhibitory to the inflammasome (Fig 6A), while not affecting TNF-α secretion (Fig 6B). Most notably, ILA alone significantly reduced caspase-1 activation to levels similar to that achieved with CM35 (Fig 6C). Interestingly, ILA did not reduce caspace-1 activation when macrophages were activated with *Δcap67* infection instead of nigericin (Fig 6D). This result suggests that ILA may participate in CM35 inhibition properties, but that it is not the sole molecule responsible for this capacity. Hence, it is possible that CM35 inhibitory properties are due to a combination of molecules. At this point we identify ILA as an immunomodulatory metabolite produced by *C. neoformans*.

**Fig 6.**
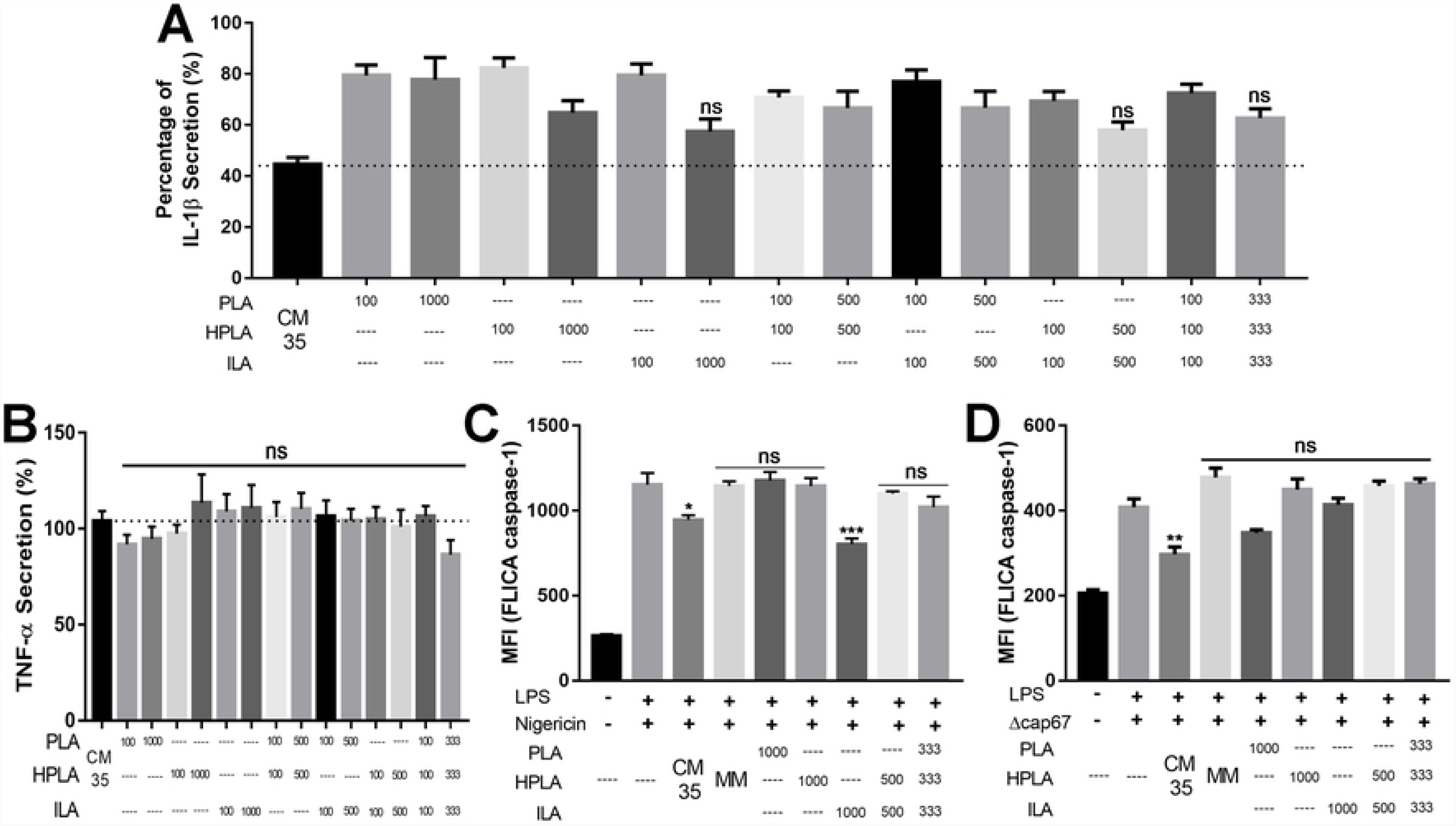
ILA is partly responsible for the inhibitory activity of CM35. A-B. Percentage of cytokine secretion, measured by IL-1β (A) or TNF-α (B) release by BMMs stimulated with LPS + Nigericin and the addition of possible inflammasome inhibitors (CM35, PLA, HPLA and ILA). Positive group LPS + Nigericin was normalized for a 100% cytokine secretion. The numbers below the bars represent the concentration of the respective metabolite in µM, except for CM35 and MM (10% v/v). The graph shows the metabolites alone, followed by double combinations and a triple combination of all metabolites. The bars represent three independent assays. C-D. BMMs stimulated with LPS (500ng/mL) and nigericin (20µM) or *Δcap67* (MOI 5:1), with or without inhibitors were analyzed for caspase-1 activation (FLICA). CM35 = Conditioned Media from C. neoformans B3501; MM = Minimal Media; PLA = DL-3-Phenyllactic acid; HPLA = DL-p-Hydroxyphenyllactic acid; ILA = DL-Indole-3-lactic acid. Statistical analysis was made utilizing One-way ANOVA, where ns: not significant; *: P ≤ 0.033; **: P ≤ 0.002; ***: P ≤ 0.001. Comparisons were made with CM35 group (black bar). ns means that the inhibition level achieved by the metabolites is similar to the inhibition achieved by CM35 (A-B). Comparisons were made with the positive control (LPS + nigericin or LPS + *Δcap67*) (C-D).

## Discussion

Despite ongoing study, many important questions regarding the processes by which fungi interact with inflammasome pathways remain unanswered. Although the major receptors involved and most of the stress signals required for inflammasome activation are known, the mechanisms by which pathogens modulate and/or evade these responses are still poorly understood. Here, we demonstrate that molecules secreted by *C. neoformans* can specifically inhibit canonical activation of an inflammasome pathway and dampen macrophage anticryptococcal activity, potentiating fungal survival and growth during the host-pathogen interaction. Furthermore, we determined that GXM did not specifically participate in this process in our model and we identified one molecule partially responsible for the effect promoted by the fungal conditioned media.

Inflammasome activation is associated with a pro-inflammatory response, mainly due to its IL-1β and IL-18 processing property. Beyond simply activating macrophages and neutrophils, both are key cytokines in the development of Th17 and Th1 polarization, respectively [44]. Moreover, since Th1/Th17 polarization is widely reported as a protective response against pathogenic fungi, NLRP3 defects are associated with a poor prognosis, ranging from severe susceptibility in invasive candidiasis [46] to milder susceptibility to cryptococcosis [52]. On the other hand, NLRP3 defects have also been associated with a better prognosis in aspergillosis associated with cystic fibrosis [63] and defects had no impact in chromoblastomycosis [50]. Hence, inflammasome activation has different effects depending on the fungal pathogen and the underlying biology of the host.

As the canonical inflammasome scaffold involves various proteins and downstream signaling, numerous steps can be inhibited to prevent scaffold formation, thus preventing the cascade at several steps with varying consequences to the cell activation. A single point of interference is sufficient to disrupt the entirety of downstream signaling and inflammasome function, from receptor activation to caspase autolysis [64]. Inflammasome activation consequences can also be partially prevented by the use of cytoprotective agents like glycine [57]. Our study demonstrated that conditioned media from *C. neoformans* strain B3501 promoted robust inhibition of IL-1β secretion and complete inhibition of LDH release, although it had less of an impact on caspase-1 activation. IL-1β can be secreted by various mechanisms, depending on the cell status, for example through membrane pores formed by Gasdermin D or through cell membrane defects during pyroptosis [65]. In this model, glycine was not able to prevent pore formation in immortalized macrophages, consequently preventing LDH release but not cytokine secretion. Studies also depicted that dying macrophages undergoing pyroptosis were the main source of IL-1β secretion in an *in vitro* model, with peritoneal macrophages exhibiting caspase-1 activation and a cytokine burst that coincided with the moment of cell death. Interestingly, caspase-1 activation inhibition prevented IL-1β secretion, but it did not alter cell death events [66,67]. Taken together, these findings show that both events can occur independently, even if they are mostly caspase-1 dependent and depend on the inflammasome stimuli, cell type and inhibition scheme utilized.

In our model, glycine and MM containing glycine dampened IL-1β and LDH release if nigericin was used as stimuli, when interacting with the cells for a long period before inflammasome activation, but not when inflammasome activation was promoted by *Δcap67* infection [57]. Another interesting feature presented in our inflammasome inhibition model was the accumulation of pro-IL-1β intracellularly in cells treated with CM35. It is well characterized that neither caspase-1 activation nor inhibition has any impact on pro-IL-1β intracellular levels in activated cells, indicating that IL-1β translation is stable regardless of whether it is subsequently cleaved [68]. One explanation for this stability of intracellular levels during caspase-1 inhibition is that pro-IL-1β, along with other inflammasome unrelated proteins, are present in the supernatant of necrotic cells undergoing NLRP3 activation [67]. On the other hand, changes in the cytokine intracellular reservoir can impact the secretion of its mature form [69]. In our work, an increase in pro-IL-1β intracellular levels occurred concomitantly with a decrease in IL-1β mature form release, which is not fully explained by the inhibition seen in caspase-1 activation. One aspect of our model is that pyroptosis is prevented in the presence of CM35, suggesting that intracellular proteins are retained during NLRP3 activation in this group. Furthermore, transcripts of *il1β* were highly expressed in the cells treated with CM35. Regarding inflammasome inhibition and pro-IL-1β, Folco et al. demonstrated that macrophages primed with LPS and treated with bafilomycin, a known inhibitor of the NLRP3 dependent receptor activation, exhibited increased intracellular levels of this cytokine [70].

The first studies depicting inflammasome activation by fungi demonstrated that morphogenesis is important to promote NLRP3 inflammasome activation, hence some morphotypes of a pathogen can induce a more robust response then others [48,71]. *C. neoformans* is also known to be a poor inflammasome activator. In this context, GXM has been considered the main factor for the yeast to evade inflammasome activation, promoting the evasion of phagocytosis and prevention of recognition by extracellular receptors [52–54].

The only well described direct modulation of the inflammasome pathway to date with a focus on fungicidal activity has been with *C. albicans* internalized hyphae in which the secretion of a candidalysin leads to cell piercing, NLRP3 inflammasome activation and host cell death via pyroptosis [72,73]. Despite the lack of further published evidence for fungal inhibition of inflammasome activation, many other intracellular pathogens, like bacteria and viruses, have well described mechanisms for NLRP3 suppression. The human pathogenic bacteria of the genus *Yersinia* express a conserved type III secretion system (T3SS) as a virulence trait, which are associated with inflammasome activation. This secretion system is responsible for the release of effector proteins called *Yersinia* outer proteins (Yops) and two Yops can inhibit inflammasome activation: YopK is related to prevention of T3SS recognition by NLR receptors [74] and YopM is related to blockage of Pyrin inflammasome activation [75]. Defects in both effector proteins lead to a more robust inflammasome activation and bacterial clearance, highlighting the importance of the inhibition promoted by the pathogen. Poxviruses produce proteins homologous to mammalian proteins responsible for inflammasome inhibition, known as pyrin domain-only proteins (POP) and serpins. These viral proteins bind to ASC or caspase domains, preventing proper inflammasome assembly and activity to promote the intracellular survival of the Poxvirus [76]. Therefore, it is not surprising to demonstrate that *C. neoformans* derived molecules can specifically inhibit inflammasome activity and it is feasible to hypothesize that other fungal driven mechanisms for inflammasome inhibition will be discovered soon.

The importance of IL-1R signaling during cryptococcosis is uncertain. While most studies have shown that IL-18 is important during infection, IL-1β itself is sometimes depicted as unnecessary for host protection [55]. However, a recent study has shown that IL-1R signaling is essential for a Th1/Th17 polarization in chronic infection model, consequentially facilitating fungal clearance by the host [56]. The data brought by these articles corroborates with the importance that is associated with the NLRP3 components during cryptococcosis [52]. In the present work, we demonstrate that macrophages treated with CM35 had a reduced capacity to control *C. neoformans* as demonstrated by a reduction in yeast cell phagocytosis, increased intracellular growth and more vomocytosis events, especially after 12 h of infection. Moreover, we did not observe any alteration in phagocytosis, fungicidal activity or vomocytosis in early time points, reinforcing the hypothesis that CM35 does not immediately affect macrophage functions. The secretion of inflammasome related cytokines is usually related to a burst, associated with cell death via pyroptosis. This burst releases a high content of pro-inflammatory IL-1 family cytokines, therefore activating other cells that did not had initially have their inflammasome pathways activated, thus promoting the maintenance of a pro-inflammatory environment [66]. On the other hand, caspase-1 inhibition severely impairs the promotion of this pro-inflammatory environment. Overall, inflammasome activation is related to a more effective response against *C. neoformans*, eliciting polarization towards Th1 and promoting an antimicrobial macrophage activation; consequently, inhibition of this pathway reduces the responsiveness of the macrophages against the fungus, as shown in this work.

Vomocytosis is here defined as a host mediated non-lytic exocytosis in which both the host cell and the fungal yeast survive. The stimulation of this event usually leads to milder disease, and is therefore considered relatively protective in one *Cryptococcus* infection model [77]. Although the mechanisms and causes of vomocytosis are still poorly understand, one of the most accepted hypotheses suggests that macrophages that undergo vomocytosis cannot control their intracellular fungal burden, which leads to these host cells releasing the fungus to the extracellular environment instead of permitting themselves to serve as a protected niche for cryptococcal replication. Alanio and colleagues demonstrated that the fate of infection is highly correlated with the intracellular growth rate of virulent strains of *C. neoformans*, showing that a high rate of replication usually increases mortality of the host, while a low rate increases the migration towards the brain [78], supporting this vomocytosis hypothesis. Our results indicate that macrophages treated with CM35 have a higher rate of vomocytosis events, suggesting that the inhibition in the inflammatory response and pyroptosis promoted by the conditioned media might also enhance vomocytosis as an alternative mechanism to expel fungal burden to mitigate against host cell death.

ILA is an aromatic metabolite derived from the tryptophan pathway. It is produced by a wide variety of organisms and microorganisms, ranging from soil bacteria to humans [61,62]. ILA production in fungi is mostly reported in endophytic and phytopathogenic species, being described as important for plant tissue colonization [79]. The tryptophan degradation pathway that leads to ILA production has an intermediate product denominated indolepyruvate, which is transformed in ILA as a result of a reduction-oxidation reaction by the NADPH dependent enzyme indole-3-lactate dehydrogenase. Although this pathway and responsible enzymes have not yet been fully annotated in *C. neoformans* nor *C. albicans*, the presence of ILA has been reported conditioned mediums from both pathogenic yeast [80]. Furthermore, the same study showed that a nitrogen source was needed for ILA production, which is supplied in our model by the presence of glycine in minimal media. The presence of a nitrogen source is very important for cryptococcal survival and growth inside the host, hence its metabolism plays a role in the expression of virulence factors [81]. In the context of pathogenesis, there is no link in the literature between ILA and fungal infections, although Zelante et al. correlated the production of tryptophan catabolites by mouse gut microbiota, among then ILA, with mucosal protection from inflammation and resistance in a candidiasis infection model, which is mediated via IL-22 production [82].

In recent years, various enzymes and proteins related to biosynthesis metabolic pathways have been depicted as important for pathogenesis in cryptococcosis infection models, especially those associated with glucose metabolism. Defects in pyruvate and hexose kinases and in acetyl-CoA production impact virulence traits of the yeast, resulting in a reduction in host mortality [83,84]. Nevertheless, our knowledge regarding aromatic metabolites derived from amino acid metabolism that are secreted by *C. neoformans* and their impact in the host during infection is severely limited. Notably, amino acid permeases are important for the protection of the yeast when challenged by environmental or host promoted stress conditions. *C. neoformans* mutants with defect in these enzymes are less virulence, highlighting the importance of amino acid uptake during infection [85]. Yeast cells deficient in a small protein allegedly involved in the citric acid cycle had a higher expression of amino acids (i.e. tryptophan) and the mutant cells were more lethal in mice compared to wild type yeast cells [86]. Another interesting aspect of the study was the demonstration that the intracellular fungal burden in macrophages infected with the mutant was only higher in the presence of exogenous NADPH, a cofactor that is essential to produce ILA from indol-3-pyruvate.

Although there is no report correlating aromatic metabolites and inflammasome inhibition, it is well known that some small molecules with similar structures are used for this effect. Glyburide is one of the most well described inhibitors for NLRP3 activation, acting on ATP-sensitive potassium channels to block potassium efflux, which prevents inflammasome activation. Glyburide is characterized as a sulfonylurea drug, but also presents aromatic hydrocarbons in its structure [87]. Interestingly, not all sulfonylurea drugs are able to prevent IL-1β secretion via inflammasome inhibition, and there are reports of sulfonylurea drugs that prevent inflammasome activation by mechanisms other than potassium efflux blockage [88].

In conclusion, this work identifies new effects mediated by *C. neoformans* wild-type conditioned media, especially in the ability of CM35 to modify the activation of the intracellular inflammasome pathway. Treating macrophages with conditioned media reduced such key functions as phagocytosis and intracellular killing in *in vitro* infection models, suggesting a possible new mechanism for fungal persistence inside the host. We also found that the aromatic metabolite ILA has a role in the inhibitory properties of CM35, while GXM and GXM-derived molecules were not involved in the activity of CM35. Nevertheless, more studies regarding *C. neoformans* conditioned media are essential to determine the detailed mechanism of action, to define where along the signaling pathway that the inflammasome is being affected, and to identify other molecules that participate in inflammasome inhibition. Thereafter, an in-depth analysis of how these molecules impact cryptococcal infection would significantly enhance our understanding of cryptococcosis and, perhaps, lead to new strategies to prevent and treat *C. neoformans* disease.

## Material and Methods

### Fungal Strains

Cryptococcal species complex strains H99 (var. grubii), B3501 (var. neoformans) and *Δcap67* (acapsular strain derived from B3501) were grown for 18 h in Sabouraud dextrose broth, rotating (120 rpm) at 30°C. Yeast cells were retrieved from culture by centrifugation (5 min, 1800 g) and washed twice in PBS before experiments.

### Conditioned medium, GXM isolation and subsequent treatments

B3501 and *Δcap67* strains were grown for 5 days in Minimal Media (MM) (glucose 15 mM, magnesium sulphate 10 mM, Monopotassium phosphate 29.4 mM, glycine 13 mM and thiamine 3 µM) rotating (120 rpm) at 30°C. [89]. Yeast cells were removed from culture by centrifugation (2x 15 min 5500 g). The supernatant was collected and filtered (0.45 µm) for complete yeast removal. The filtrate was lyophilized and suspended in deionized water, with a tenfold concentration. The products obtained from the B3501 and *Δcap67* strains were denominated Conditioned Media 35 (CM35) and Conditioned Media CAP (CMCAP), respectively [90]. CM35 was treated and/or fractioned for subsequent experiments. The size fractions were obtained utilizing an ultrafiltration Amicon system (Millipore), with filtration membranes varying from 100 to 1 kDa. In between fractions (e.g.: 100 kDa> CM35 > 10 kDa) were also obtained, by recovering molecules retained in the filtration membrane. Polarity fractions were obtained by Blight-Dyer technique. Additionally, CM35 was treated by autoclaving (20 min at 123°C) and with the following proteases (24 h at 37°C): trypsin, thermolysin and pronase. CM35 was also processed to remove GXM using a GXM-capture ELISA, as described by Rodrigues et. al. [60]. Briefly, an ELISA high-binding plate was coated with mAb 18B7 (a monoclonal antibody (Ab) specific for GXM [91] for 2 h at RT, preceded by a blocking step with 1% BSA solution for 1 h at RT. Finally, conditioned media or minimal media were added to the wells for additional 2 h and recovered at the end. The bound GXM was recovered by elution with Tris-Glycine (pH 7.4) buffer. Yeast capsular polysaccharides from B3501 were harvested [92] and kindly supplied by Julie M. Wolf (Albert Einstein College of Medicine). Exo-polysaccharides were obtained by the collection of a viscous layer in the 10kDa membrane during CM35 ultrafiltration, as described [93].

### Polysaccharide (PS) attachment and immunofluorescence

Polysaccharide (PS) attachment to *Δcap67* cell wall was performed as described [94]. *Δcap67* cells were incubated in yeast-free B3501 conditioned medium (grown in minimal medium for 4 days) overnight at 37°C. B3501, *Δcap67* and *Δcap67*-PS were fixed with 4% paraformaldehyde for 30 minutes at room temperature and incubated with PBS + 1% BSA for 1 h at room temperature. Yeast cells were then incubated with 0.01% Uvitex2B (a chitin marker; Polysciences) and mAb 18B7 (10 µg/ml) for 30 minutes followed by incubation with Alexa-Fluor 546 anti-mouse IgG1 (5 µg/ml; Invitrogen) for 30 minutes at 37°C. Cells were then suspended in an anti-fading agent, mounted on glass slides and analyzed with a confocal microscope (Leica TCS SP5).

### Ethics statement

All experimental procedures were approved by the Animal Ethics Committee of the University of Brasilia (UnBDoc number 55924/2016) and conducted according to the Brazilian Council for the Control of Animal Experimentation (CONCEA) guidelines.

### Generation of bone marrow-derived macrophages (BMMs) and dendritic cells (BMDCs)

Bone marrow cells were obtained from C57BL/6 isogenic mice (8–10 weeks old), as previously described [49]. Briefly, femurs and tibias were flushed with RPMI-1640 to harvest the bone marrow cells. Cells were submitted to erythrocyte lysis under a tris-buffered ammonium chloride solution, seeded (2 × 10^5^ cells/ml) and cultured for 8 days at 37°C in non-tissue culture-treated Petri dishes in 10 ml/ dish of RPMI-1640 medium that contained 50 mM 2-mercapto-ethanol. The medium was supplemented with 20 ng/ml murine granulocyte-macrophage colony-stimulating factor (GM-CSF, Peprotech) or 30% conditioned medium from macrophage colony-stimulating factor-secreting L929 fibroblasts (M-CSF) to obtain BMDCs and BMMs, respectively. On the third day, additional 10 mls of complete medium that contained differentiation-inducing cytokines were added. Half the medium was exchanged on the sixth day of culturing the BMDCs. On the eight day, non- and loosely adherent BMDCs or firmly adherent BMDMs/BMMs were stripped with TrypLE™ Express (Gibco), harvested and plated in RPMI-1640 medium supplemented with Fetal Bovine Serum (FBS) and gentamicin.

### Murine cell cultures interaction with conditioned media

BMMs or BMDCs (1 × 10^6^/mL) were incubated at 37°C in a humidified 5% CO^2^ atmosphere. Cells were stimulated with lipopolysaccharide (LPS; 500 ng/mL for 4h – Sigma-Aldrich), providing the first signal for inflammasome activation. Additionally, cells were incubated for 18h with or without potential inhibitors: CM35 (and its fractions), CMCAP, Minimal Medium, Glycine (Sigma-Aldrich) Glucuronoxylomannan (GXM) and aromatic metabolites (Sigma) 3-Phenyllactic acid (PLA); DL-p-Hydroxyphenyllactic acid (HPLA) and DL-Indole-3-lactic acid (ILA), alone or in combination. Thereafter, cells were treated with nigericin (20 µM for 40 or 90 minutes – Invivogen), providing the second signal for inflammasome activation. Alternatively, cells stimulated with LPS were infected with the fungal strains opsonized with mAb 18B7 (a kind gift from A. Casadevall, Johns Hopkins Bloomberg School of Public Health) [91]. Controls included conditions without LPS or nigericin.

### Cytokines quantification by enzyme-linked immunosorbent assay (ELISA) and LDH detection

The cell-free supernatants of the BMM and BMDC cultures were harvested for measurements of IL-1β and tumor necrosis factor (TNF)-α (Ready-Set-Go! Kit – eBioscience) concentrations using ELISA. The determination of intracellular pro-IL-1β was performed with the cell lysates (Ready-Set-Go! Kit – eBioscience). The data were expressed as pg/ml ± the standard deviation (SD) of two to three independent experiments, which were conducted in triplicate.

The cell-free supernatants of the BMM cultures were harvested to quantify LDH release, as a cell death marker (CytoTox 96^®^ Non-Radioactive Cytotoxicity Assay – Promega) after treatment with nigericin for 90 minutes. The data were expressed as percentage ± the standard deviation (SD) of two to three independent experiments, which were conducted in triplicate, considering cells without any treatment as 0% and cells treated with 15% DMSO as 100% of cell death.

### Active caspase-1 detection by flow cytometry

BMMs challenged with *C. neoformans* strains (MOI – 5:1) opsonized with mAb 18B7 were harvested from the tissue culture well plates with a dissociation agent (Tryple Express – Thermo-Fischer Scientific). 5 × 10^5^ were collected per group before staining. Staining for caspase-1 (FAM-FLICA™ Caspase-1 Assay Kit – Immunochemistry) was performed according to the manufacturer’s instructions. The cells were then analyzed in a flow cytometer (FACSVerse – BD Biosciences) with the FITC filter channel (FL-1) and data were processed (FlowJo × – LLC).

### cDNA synthesis and real-time RT-PCR

Total RNA from the cultured BMMs was extracted using TRIzol reagent (Invitrogen) and cDNA synthesis was performed using a high capacity RNA-to-cDNA kit (Applied Biosystems), according to manufacturer’s instructions. qRT-PCR was performed using SyBr Green Master Mix and StepOne real-time PCR system (Applied Biosystems). The primer sequences were as follows: IL-1β forward, 5’- GTGTGTGACGTTCCCATTAGACA-3’; IL-1β reverse, 5’-CAGCACGAGGCTTTTTTGTTG-3’; Nf-κB forward, 5’-AGCCAGCTTCCGTGTTTGTT-3’; nuclear factor kappa-light-chain-enhancer of activated B cells (Nf-κB) reverse, 5’-AGGGTTTCGGTTCACTAGTTTCC-3’; GAPDH forward, 5’- TGAAGCAGGCATCTGAGGG-3’; Glyceraldehyde 3-phosphate dehydrogenase (GAPDH) reverse, 5’- CGAAGGTGGAAGAGTGGGAG-3’; IL-1β or Nf-κB to GAPDH relative expression was calculated using the 2^(-ct)^ method and normalized to the level of unstimulated BMMs.

### Vomocytosis scoring by flow cytometry

BMMs challenged with stained H99 with Calcofluor White (10 µg/mL, 10 min – Sigma-Aldrich) were treated concomitantly with CM35 or left untreated. After 2 h of interaction, all wells were gently washed three times with warm PBS. The last wash from the 2 h time point wells was collected and the cells were lysed. RPMI with fluconazole (10 µg/mL – Sigma-Aldrich) was added in the other wells and the cells were treated again with CM35 or left untreated. The supernatant and cell lysate were collected in determined time points (6, 12 and 24 h). Right after the collection, both supernatant and lysate samples were centrifuged (2.000xg, 5 min) for yeast recovery and fixated in PFA 4% in PBS until flow cytometry analysis. Samples were accessed by flow cytometry (FACS Fortessa – BD Biosciences), at low speed acquisition for a 4 minutes period. Parent yeast cells were detected using a high calcofluor fluorescence gate. Using the specification provided by the manufacturer, the number of yeast/mL was calculated.

### Co-incubation assays (phagocytic index, fungicidal activity and transwell assays)

BMMs were challenged with H99 opsonized with mAb 18B7 (MOI – 2:1) and the phagocytosis rates and fungal burdens were analyzed at 37°C. For the phagocytosis assay, BMMs were primed with LPS (500 ng/mL) for 4 h followed by treatment overnight with conditioned media. Controls were left untreated. After this period the cells were challenged with opsonized H99. After 2 h, the wells were gently washed three times with warm PBS and the remaining co-culture was fixed and stained with modified Wright-Giemsa stain. The phagocytic index was calculated by multiplying the number of macrophages engaging phagocytosis and the number of yeasts phagocyted per 100 macrophages analyzed. For the yeast intracellular growth assay, BMMs primed with LPS (500ng/mL) or not were challenged with opsonized H99 and concomitantly treated with conditioned medias or left untreated. After 2 h, all the wells were gently washed three times with warm PBS to remove extracellular yeast. Right after the washes as well as 24 h later, macrophages were lysed with 0.05% SDS and intracellular yeasts recovered and measured by colony forming units (CFU) by plating on Sabouraud dextrose agar. The intracellular growth rate was determined by dividing 2h CFU by 24h CFU. For the transwell assay, BMMs were seeded in both upper and lower chambers and primed with LPS (500ng/mL). The BMMs in the lower chamber were challenged with opsonized *Cryptococcus* strains (MOI 5:1) or left uninfected, while the BMMs in the upper chamber were not challenged. After 24 h at 37°C, macrophages from the upper chambers were challenged with opsonized GFP-expressing H99 (MOI 2:1) and lysed with 0.05% SDS at 24 h post-infection. Alternatively, the lower chambers contained only yeast cells from *Cryptococcus* strains in a media with LPS (500ng/mL) throughout the assay. The recovered intracellular yeast cells were quantified using flow cytometry.

### Detection of GXM internalization using fluorescence microscopy

BMMs primed with LPS were treated with CMCAP or CM35, and their fractioned derivatives, as described earlier. BMMs were plated in glass inserts and after 18 h of interaction with conditioned media, cells were permeabilized and fixed with cold methanol (Vetec) for 10 minutes. After consecutive washes, cells were blocked for one h with a 10% FBS solution and stained for GXM with mAb 18B7 (10µg/mL) as a primary Ab and “Alexa Fluor® 633 Goat Anti-Mouse IgG” (Life Technologies) as a secondary Ab. Cells were also stained with a DAPI solution (Life Technologies). The glass inserts were then recovered and GXM content internalized by the BMMs were observed under a confocal microscope (Leica TCS SP5).

### Gas chromatography-mass spectrometry analysis

Metabolites from conditioned culture media were derivatized as described [95]. Ketone groups of metabolites were derivatized by adding with 20 μL of 30 mg/mL methoxyamine in pyridine (Sigma-Aldrich) and incubating at 37°C for 90 minutes with shaking (1000 RPM). Then, amine, hydroxyl and carboxyl groups were modified with 80 μL of N-methyl-N-(trimethylsilyl) trifluoroacetamide (MSTFA) (Sigma-Aldrich) with 1% trimethyl-chlorosilane (TMCS) (Sigma-Aldrich) by shaking (1000 rpm) at 37°C for 30 minutes. Derivatized samples were analyzed in an Agilent GC 7890A using a HP-5MS column (30 m × 0.25 mm × 0.25 μm; Agilent Technologies) coupled with a single quadrupole MSD 5975C (Agilent Technologies). Samples were injected in the splitless mode with the port temperature set at 250 °C and oven temperature equilibrated at 60°C. The oven temperature was kept at 60°C for 1 minute, then raised to 325°C at a rate of 10°C/minute, and finally finished with 5 minutes hold at 325°C. A fatty acid methylesther (FAME) standard mix was analyzed with each batch for subsequent retention time calibration purposes. Collected data files were processed with Metabolite Detector [96], by calibrating retention indices (RI) based on FAME standards, and deconvoluting and chromatographically aligning features. Metabolites were identified by matching spectral features and retention indices against a PNNL augmented version of the FiehnLib library containing more than 900 metabolites [97]. Unidentified metabolites were also searched against the NIST14 GC-MS library by comparing spectral features only. All identifications were manually validated. Extracted metabolite intensities were subjected to multivariate data analysis (MVDA) using MetaboAnalyst [98]. The data were median normalized and log transformed followed by principal component, hierarchical cluster, and heatmap analysis to identify natural clustering within the data.

### Statistical Analysis

Statistical analysis was conducted using GraphPad Prism v.7.0 software. Data were analyzed by one-way ANOVA followed by Tukey’s post hoc test. *p*-Values of less than 0.05 were considered significant. The metabolomics assay was analyzed by t-test, PCA and OPLS, the metabolite figure was generated from an excel spreadsheet that was extracted from a heatmap using the t-test values. For normalization, the data were median centered and log transformed.

## Acknowledgements

We thank Stephan Oliveira Machado (University of Brasília) and Guilaume E. Desanti (University of Birmingham) for technical assistance. We thank Julie M. Wolf (Albert Einstein College of Medicine) for the capsule polysaccharides. We thank Dario Zamboni (University of São Paulo) and Kelly Magalhães (University of Brasília) for discussions.

## Supporting Information

**S1 Fig. CM35, but not CMCAP or Minimal Media is able to reduce IL-1β secretion.** Secretion of IL-1β and TNF-α were measured from BMDMs stimulated with LPS (500ng/mL) and/or nigericin (20µM), with or without possible inhibitor (10% v/v) overnight (18h) (A and B). Alternatively, BMMs were treated with possible inhibitors overnight (18h) previously to stimulation with LPS (500ng/mL) for 4 h (C). Supernatants were collected after stimulus and cytokines measured by ELISA technique. CM35 = Conditioned Media from B3501; CMCAP = Conditioned Media from *Δcap67*; MM = Minimal Media. Statistical analysis was performed utilizing One-way ANOVA, where ns: not significant; *: P ≤ 0.033; ***: P ≤ 0.001.

**S2 Fig. CMs induce IL-1β transcription in activated macrophages.** Transcript levels of IL-1β (A) and Nf-κB (B) from cDNA extracted from BMMs stimulated with LPS (500ng/mL) and/or nigericin (20µM), with or without potential inhibitor (10% v/v) overnight (18h). IL-1β or Nf-κB to GAPDH relative expression was calculated using the 2^(-ct)^ method and normalized to the level of unstimulated BMMs. Statistical analysis was achieved by One-way ANOVA, where ns: not significant; *: P ≤ 0.033; ***: P ≤ 0.001. Comparisons were made with the LPS + nigericin group.

**S3 Fig. Transwell assay membrane integrity analysis.** Flow cytometry analysis (24h post infection) from intracellular yeast cells in BMMs infected with H99 expressing GFP (2:1), in the upper chamber of a transwell apparatus. In the lower chamber, BMMs were stimulated with LPS (500ng/mL) alone (A) or together with B3501 (B) or *Δcap67* (C).

**S4 Fig. GXM detection in CM samples.** A-C. BMMs interacting with inhibitors CM35 (A), CM35 <1kDa (B) and CMCAP (C) D-F. BMMs interacting with CM35 inhibitor before GXM depletion treatment by capture ELISA (D), after treatment (E) and enriched with GXM eluted from the ELISA plate (F). Cells were stained for nucleus (blue) and GXM (red). Images were taken in a confocal fluorescence microscope.

**S5 Fig. Mass spectrometry comparative CM analysis.** A – Heat map based on extracted metabolite significant intensities from CM35 and CMCAP. B-D Comparative data from each aromatic metabolite detected in the analysis.

